# Thermal conditions and habitat size explain elevational range shift of the northern pika

**DOI:** 10.64898/2026.08.06.743106

**Authors:** Tomoki Sakiyama, Jorge García Molinos

## Abstract

**Aim:** Species in mountain ecosystems often experience upslope distribution shifts in response to climate change. However, elevational range dynamics exhibit substantial complexity around this general trend, with varying magnitude and direction of shifts observed among studies. We tested whether factors other than temperature, such as habitat size and land use, are also responsible for observed elevational shifts in a cold adapted species.

**Location:** Hokkaido, Japan

**Methods:** We resurveyed 61 historical (1963-2007) occurrence sites covering a wide elevational range (60–2,210 m) within the distribution range of the northern pika (*Ochotona hyperborea*). We assessed the existence of elevational shifts using quantile regression and the relative importance of temperature, habitat size, and land use on the shift using occupancy analysis.

**Results:** We detected presence of the northern pika at 41 sites, suggesting extirpations at 20 sites (32.8%) across a wide elevational range of 60–1,550 m. The concentration of extirpation sites at low to mid elevations resulted in a significant upslope shift of the distribution centroid, although the shift was nonsignificant at lower and upper portions of the range. The occupancy analysis revealed a negative effect of long-term mean of summer maximum temperature and a stronger positive effect of habitat size.

**Conclusions:** This highlights the susceptibility of the northern pika to heat stress and the importance of larger habitats and thus habitat heterogeneity for persistence of local populations. Given the possibility that the observed shift represents a precursor of elevational contraction, continuous monitoring of the local populations is highly needed to evaluate their long-term viability.

## INTRODUCTION

Climate change is one of the main threats to global biodiversity. Accumulating evidence suggests that numerous species are shifting their distributional ranges in response (Parmesan and Yohe 2003, Chen et al. 2011, Lenoir et al. 2020), with predicted consequences for interacting species as well as ecosystem services and functions (Pecl et al. 2017). With future climate projected to change continuously, understanding the drivers of range shifts is critical for informing conservation and management decisions (Bonebrake et al., 2018; Dawson et al., 2011).

Changes in species distribution are frequently evaluated based on resurveys, where locations surveyed for species presence or absence in the past are revisited to detect changes in occurrence over time (Tingley and Beissinger 2009, Verheyen et al. 2017). In mountain ecosystems, resurveys conducted along elevational gradients have revealed upslope shifts (Moritz et al. 2008), including expansion of upper elevational limits (Rowe et al. 2009, Tingley et al. 2012), contraction of lower limits (Wilson et al. 2005), and upward shifts of elevational optima (Lenoir et al. 2008, Bergamini et al. 2009). Because temperature declines linearly with elevation and strongly constrains the physiological performance of many species, these observed patterns are generally consistent with *a prirori* expectations under climate warming. However, elevational range dynamics exhibit substantial complexity around this general trend (Rubenstein et al. 2023). The magnitude of observed shifts varies widely among both individual species (Moritz et al. 2008) and broader taxa (Lenoir et al. 2020), with absence of detectable shifts frequently reported (Chen et al. 2011, 2025). Moreover, downslope shifts have also been documented (Crimmins et al. 2011, Bhatta et al. 2018). Variability in responses has been observed even within species at the population level (Rowe et al. 2015) and across different positions within species’ ranges (Tingley et al. 2012). Together, these patterns suggest that temperature alone is insufficient to fully explain observed changes in species distributions (Rapacciuolo et al. 2014, Chen et al. 2024). Despite its key importance on physiological processes and frequent correlation with other abiotic parameters, species range shifts ultimately result from the complex interplay of multiple abiotic and biotic factors (Lawlor et al. 2024). Accordingly, some resurvey studies have incorporated additional aspects of climate change, such as changes in precipitation conditions, to better explain observed range shifts (Crimmins et al. 2011). Nevertheless, the effects of non-climatic factors remain comparatively understudied (Rapacciuolo et al. 2014), despite predictions that, where habitat quality is governed by non-climatic conditions, climate warming may lead local populations to extinctions more frequently in low-quality habitats than in high-quality habitats (Brook et al. 2008, Mantyka-Pringle et al. 2015). Given that species are simultaneously exposed to multiple environmental conditions, incorporating non-climatic drivers into resurvey analyses is essential for understanding range dynamics (Lenoir and Svenning 2015, Chen et al. 2025). For instance, anthropogenic factors, such as human land use, may influence species persistence over time through alteration of natural habitats (Mantyka-Pringle et al. 2015, Sokolova et al. 2024). Moreover, habitat patch size is important for local population viability, with larger habitats typically having greater environmental heterogeneity and carrying capacity, hence sustaining larger population sizes and leading to higher persistence (Griffen and Drake 2008).

Pikas are small lagomorphs occurring in mountain ecosystems and are perceived as vulnerable to climate change (Wang et al. 2020). Often recognized as alpine specialists, some pika species are nonetheless also known to occur at lower elevations (Varner and Dearing 2014). These populations are likely supported by the rocky landforms they inhabit, such as talus slopes and lava flows (Gliwicz et al. 2005, Rodhouse et al. 2010, Smith et al. 2018), which buffer ambient air temperatures by locally generating cool and thermally stable conditions (Varner and Dearing 2014, Millar et al. 2016, Sakiyama et al. 2021) reducing physiological stress (Wilkening et al. 2015). Consequently, such complex topography is expected to mitigate effects of climate warming and contribute to the viability of local populations (Morelli et al. 2016). Nevertheless, resurveys of pikas have indicated substantial distributional changes, with lower-elevation populations becoming extirpated under warming conditions (Beever et al., 2003, 2011; but see Millar & Smith, 2022). However, most of these studies have primarily been conducted on the American pika (*Ochotona princeps*), with a focus on climatic drivers, while studies investigating responses of other pika species to both climatic and non-climatic drivers remain scarce.

The northern pika (*Ochotona hyperborea*) occurs across northeastern Eurasia and has the largest distribution range among all pika species. The southern marginal populations are found on Hokkaido Island, Japan, which are presumably the most vulnerable to climate change given their southern location and the strong influence of temperature on their distribution (Sakiyama et al. 2021, Sakiyama and García Molinos 2024). The entire metapopulation is designated as near threatened in the Japanese Red List due to limited extent of suitable habitat (Ministry of Environment 2014, 2020). However, despite existing extensive historical records of its distribution from the 1960s to the 2000s (Onoyama and Miyazaki 1991, Kawabe 2008) and evidence of accelerating warming in Hokkaido (Higashino and Stefan 2014), resurveys on the northern pika have not been conducted to assess the impacts of climate change on its distribution. Further, the geographical range of the northern pika in Hokkaido encompasses great environmental heterogeneity, spanning an elevational gradient from near sea level to the highest elevations on the island. Although a recent study revealed the negative influence of human land use on occupancy (Sakiyama and García Molinos 2024), its presence has been reported both inside and outside protected areas in the past, providing an opportunity to investigate the combined effects of climate change and anthropogenic pressures on changes in historical occupancy at varying intensities. To do so, here we assess contemporary changes in the distribution of the northern pika in Hokkaido by relating patterns of persistence at 61 historical (1963-2007) sites spanning a wide elevational gradient (60–2,210 m), to the combined effects of temperature, habitat size, and anthropogenic factors (i.e., land use). To the best of our knowledge, this study represents the first resurvey-based assessment of contemporary range shifts in the northern pika, providing a much-needed update on the distribution of the northern pika in Hokkaido.

## MATERIALS AND METHODS

### Study area

Within Hokkaido, the northernmost island of Japan, the northern pika occurs in various mountain ranges, exhibiting a scattered spatial distribution (Onoyama and Miyazaki 1991). To compare its historical and contemporary distribution, we initially selected our study sites based on previous studies reporting the presence of the northern pika, thus allowing inference for persistence and extirpations at those sites (Tingley and Beissinger 2009). We could not include historically absent sites because such cases were often not recorded in previous studies. Given the northern pika distribution is reported from various regions in central Hokkaido, an ideal design would have included regional replicates of the elevational gradient (McCain and Grytnes 2010). However, this was not feasible because mountain ranges differed substantially in their elevational ranges among regions, and because occupied sites were typically reported from a limited portion of each region. Consequently, our study was based on a single elevational gradient constructed by aggregating sites across regions in Hokkaido. The initial selection process followed these criteria: (1) historical survey period was before 2010, (2) the site location was indicated by a map and/or a detailed description (e.g., 1,738 m point in the Mt. Tomuraushi trail), and (3) the survey methods were described clearly. Although a total of 362 sites initially satisfied these criteria, given time and logistic constraints with the resurveys, we further narrowed the selection of our study sites to best represent the reported historical distribution of the species by conducting a principal component analysis focusing on elevational, thermal, and anthropogenic gradients (i.e., land use, protection status) (Fig. S1) and considering site accessibility. In this process, we also considered sites surveyed from a recent study on northern pika occupancy in Hokkaido that have also been surveyed historically (Sakiyama and García Molinos 2024, 2026). This resulted in a final set of 61 study sites, including 45 sites newly surveyed in this study for contemporary status, that were historically reported in 8 studies from 6 regions with mean, minimum, and maximum number of sites per region of 9.6, 4, and 16 sites, respectively (Figs. 1, S1). The oldest, median, most recent record of presence were from 1963, 1989, and 2007, respectively. Although site elevations were often reported in previous studies, we used the 10 m-resolution digital elevation model provided by the Geospatial Information Authority of Japan (2023) to ensure consistent comparisons across sites. Based on this dataset, our final set of study sites spanned the elevational range of 60–2,210 m, including both the lowest and the highest elevation documented in the records. Roughly, sites with elevations of < 1,200 m, 1,200–1,600 m, and >1,600 m were characterized by forest vegetation, subalpine vegetation (forest-alpine ecotone), and alpine vegetation, respectively. A total of forty-five sites (73.8% of the total) were within existing protected areas, such as the Daisetsuzan National Park (DNP), the Hidaka-Sanmyaku Erimo Tokachi National Park (HSETNP), and the Furano-Ashibetsu Prefectural Natural Park (FAPNP). Chronologically, DNP and FAPNP were established before the historical surveys, whereas the predecessor of HSETNP was established as a Quasi-National Park in 1981, overlapping with the period of the historical surveys. Because this may introduce a confounding effect of changes in protection status over time, we instead accounted for anthropogenic effects directly by assessing the presence of human land use (see *Environmental factors* for details). All sites above 920 m were located within protected areas, but this was unavoidable because of the strong elevational bias of protected area designation toward higher elevations.

**Fig. 1.**
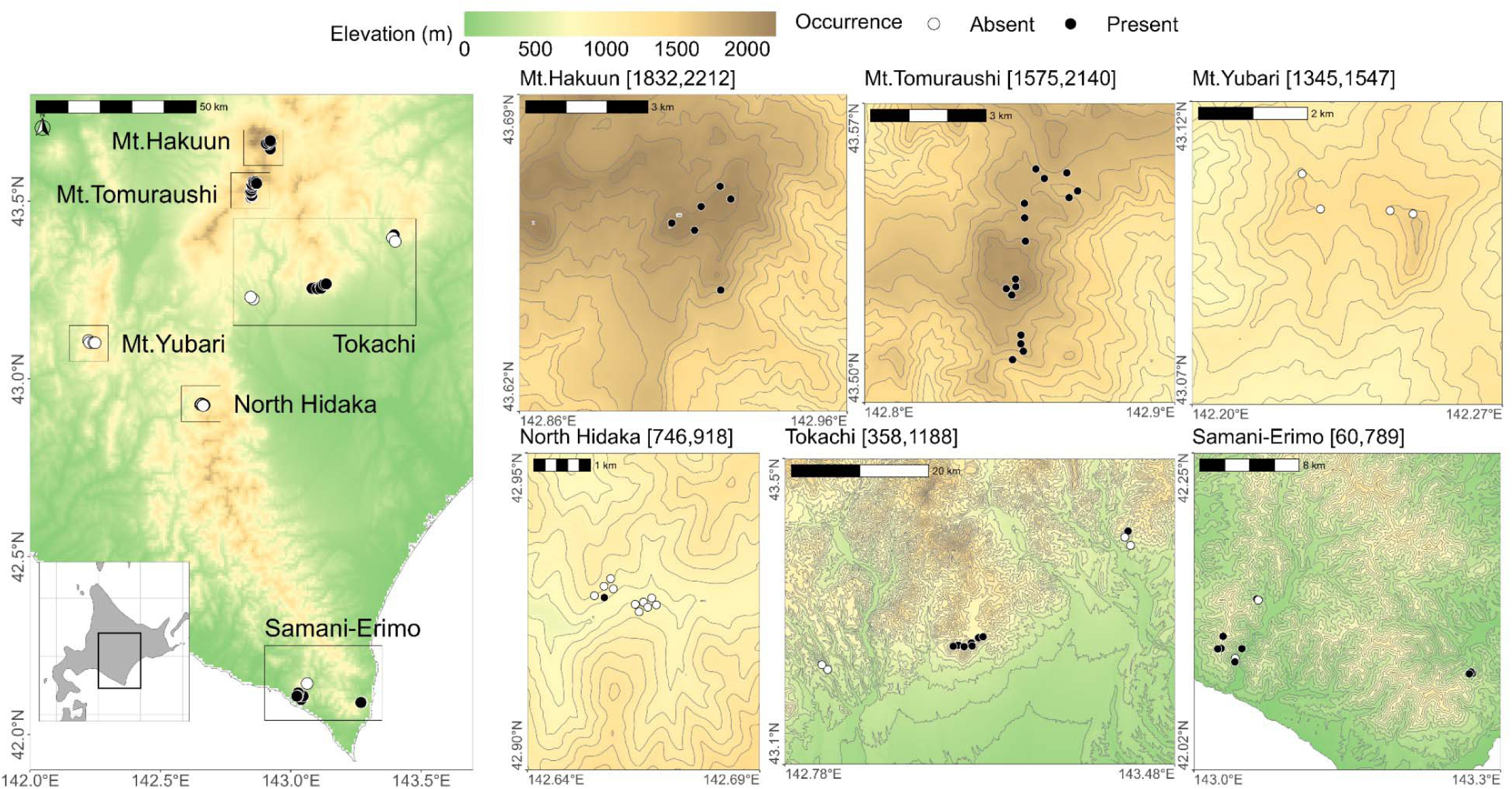
Map of the study area showing the six resurvey regions and the observed presence (black points) or absence (white points) of the northern pika at each study site. Background colors represent elevation, and values shown after region name denote the elevational range within each region.

### Resurvey

We conducted the resurvey of the northern pika during the summers of 2021 to 2023 by assessing its presence or absence at the selected sites. To account for the large environmental variability among sites, we adopted a more flexible modification of the playback method described in Sakiyama and Garcia Molinos (2023), which consisted of broadcasting a pre-recorded northern pika vocalization from a loudspeaker and listening for responding vocalizations to determine the presence (response vocalization) or absence (no response vocalization) of the pika at each site. In a single survey, playback was implemented multiple times during the approach to, the stay at, and the leave from each site until a response was heard (in cases of presence) or until the site and its periphery were fully surveyed (in cases of absence). The waiting time for vocalization responses after each playback was three minutes, which has been found to be long enough to detect most of the responding vocalizations from the northern pika (Sakiyama and García Molinos 2023). To reduce the possibility of false-negative detections, surveys were conducted two or three times at each site over the three-year period; however, some sites where northern pikas were detected during the first survey were surveyed only once. In contrast, for sites surveyed in the previous study, only a single playback was conducted in a single survey with a waiting time of five minutes, and a maximum of nine surveys were conducted over the three-year period (Sakiyama and García Molinos 2024). Although differences in the number of playbacks and surveys could potentially influence detection probability through variation in survey effort, a positive relationship between them was not observed (Fig. S2), supporting the integration of results from different survey schemes in the present analysis. The contemporary state of each local population was considered present if at least one detection occurred across multiple surveys and as absent if presence was not detected in any survey.

The exact locations of some historical sites (*n* = 8), located in the North Hidaka (87.5%) and Mt. Yubari (12.5%) regions, were difficult to determine based on the map reported in previous studies since we could not detect a typical northern pika habitat in the field (Fig. S3). In all of these sites, originally reported in Haga et al. (1979) and Kojima & Kawamichi (2001), rocks were found at the reported locations but they were embedded in soil, litter, and dense vegetation, resulting in absence or limited number of rock interstices (Fig. S3a, c, d). However, given that other sites from the same reports located in proximity to these sites exhibited characteristics of a northern pika habitat (i.e., rocky landforms with numerous interstices) including a detected presence in the North Hidaka region (Fig. S3b), we assumed that they were reliable in cartographic quality and survey results. Therefore, we kept these sites in the analysis as current absent sites. Nonetheless, to reflect the uncertainty in site location, we identified them as “low-confident sites” and considered their effect on the results by conducting sensitivity analyses both including and excluding them.

### Environmental factors

To assess the environmental factors affecting the current northern pika distribution, we considered the effects of thermal condition, habitat size, land use at each site. For thermal condition, we focused on the mean and temporal change of annual mean temperatures and summer (July-August) maximum temperatures over the last 44 years (1980–2023). Annual and summer maximum conditions were respectively used to characterize the general thermal environment and to capture potential summer heat stress considering the weak heat tolerance of pikas (Smith 1974). The long-term mean was used to reflect thermal conditions generally driven by elevation, whereas temporal change was used to characterize the speed of thermal changes ongoing at each site (i.e., warming rate) (Fig. S4). We hypothesized that extirpations occur at sites with higher mean temperatures and larger thermal changes (i.e., greater and more rapid thermal exposure). To derive these variables, we used the Agro-Meteorological Grid Square Data provided by the National Agriculture and Food Research Organization (Ohno et al. 2016, Murakami 2026); an interpolated climate data based on weather station observations. Mean daily temperature data were first used to calculate annual mean temperatures, which were then used to compute the long-term mean and temporal change over the 44-year period. Temporal change was quantified as the slope of a linear regression of yearly mean temperature against year. Similarly, daily maximum temperatures for July and August were used to calculate annual means of summer maximum daily temperature, from which the long-term mean and temporal change were derived. Although the original data had a spatial resolution of approximately 1 km, calculated annual mean values were downscaled to 100 m resolution using regression kriging implemented in the *gstat* R package (Pebesma 2004, Gräler et al. 2016), with elevation data from the Geospatial Information Authority of Japan (2023) resampled to 100 m resolution. We then extracted raster cell values of each temperature metric within a 200-m radius of each site and used the spatial average for each of these variables. This buffer distance was chosen to reflect the reported home range size of the northern pika (1,822–11,530 m^2^; Onoyama et al., 1991). The starting year was set to 1980 due to the limited availability of high-resolution climate data prior to this year.

Habitat size was defined by calculating the area (width × length) covered with rocks and interstices measured in the field. For some sites characterized by strict protection schemes and/or limited navigability due to steep, complex terrain (*n* = 23), we quantified the habitat size using aerial images in QGIS (QGIS Development Team 2023). In these cases, the maximum habitat size was capped at 10,000 m² (100 m × 100 m) to align with the area generally covered in field surveys. However, this aerial image approach still failed to quantify habitat size for sites along steep ridgelines with forested vegetation (*n* = 3), although we successfully detected northern pika presence at these sites from the adjacent trail using playback. Thus, these sites were included to compare historical and contemporary states but excluded from the final analysis of factors underlying range shift (see “Statistical analysis”). The habitat size of the low-confident sites described before was recorded as zero since all these sites lacked rock interstices.

We evaluated the presence of human land use in the vicinity to each site as an index of anthropogenic disturbance, which could influence persistence of the northern pika negatively (Sakiyama and García Molinos 2024). We inspected the presence of human land use at each site using aerial imagery from the late 2010s provided by the Geospatial Information Authority of Japan (Accessed March 7th, 2024) by identifying signs of land conversion within a 200-m radius of each site. Although various activities such as dam and road construction as well as forest plantations were detected, and these land conversions may have occurred before and/or after the historical surveys at varying intensities, we did not distinguish either the type, timing, or intensity of conversion due to the limited availability and spatial resolution of aerial imagery covering the period of historical surveys. Instead, we simply characterized human disturbance to represent contemporary land use conditions at our study sites and treated it as a binary variable (presence or absence) in the analysis. While our study sites were selected from both within and outside protected areas, this index was strongly associated with site protection status (χ² = 16.0, df = 1, *p* < 0.001).

### Statistical analysis

To test elevational shifts from past to present, we compared elevations of occupied sites at different positions within the elevational distribution using quantile regression (Koenker et al. 2017). This method estimates conditional quantiles of the response variable and enables assessing the magnitude and direction of shifts across the elevational distribution without assuming normality or constant variance. We fitted models for the 10th, 50th (median), and 90th percentiles of elevations with survey period (past vs. current) as a categorical predictor to assess shifts in the lower, central, and upper portions of the elevational distribution using the *quantreg* R package (Koenker 2025). Statistical significance was assessed using case-resampling bootstrap standard errors with 1,000 replications. We also compared the mean elevations between survey periods using a two-sided *t*-test. To account for uncertainty associated with low-confidence sites, analyses were conducted both including these sites (*n* = 61) and excluding them (*n* = 53).

We used a regression model framework to examine the effects of environmental factors on contemporary occupancy. Because study sites were constrained to previously occupied sites in our study, contemporary presence and absence reflect persistence and extirpation, respectively. Thus, this analysis is essentially equal to assessing factors that maintained persistence and drove extirpation. We included thermal conditions (mean and temporal change), habitat size, and land use occurrence as predictor variables. Although thermal variables representing both annual and summer conditions were considered, preliminary analyses focusing on mean conditions indicated better model fit when using summer maximum temperature (Table S1). Hence, summer thermal conditions were used in the global model. Prior to modeling, all predictor variables were standardized (mean = 0, standard deviation = 1) to facilitate comparison of their relative importance. Correlations among predictor variables were assessed using Pearson’s correlation coefficient for continuous variables (i.e., temperature and habitat size) (Fig. S5) and point-biserial correlation coefficients for comparisons between continuous and categorical variables (i.e., land use) (Table S2). This analysis revealed a strong correlation between the mean and temporal change of summer maximum temperature. Based on interpretability, the mean summer maximum temperature was retained in the model. All other pairwise correlations were below the commonly used threshold for strong collinearity (|r| > 0.7). We initially considered using a mixed-effects model including study region as a random effect to account for potential spatial autocorrelation in the residuals. However, this produced a near-zero variance estimate for the random effect, resulting in a singular fit. This situation can arise when fixed effects capture much of the spatial structure in the data, which is plausible in our case because thermal conditions at large explain the regional differences. Therefore, we fitted a generalized linear model (GLM) with a binomial error distribution and a logit link function using the *lme4* R package. To identify the parsimonious model, we constructed a set of candidate models including all possible combinations of the predictor variables (Table 1) and ranked them using Akaike’s Information Criterion corrected for small sample sizes (AICc) with the *MuMIn* R package (Bartoń 2010). The model with the lowest AICc was considered the best supported, while models with ΔAICc < 4 were regarded as having comparable support (Burnham and Anderson 2002). In the latter case, we calculated model-averaged parameter estimates based on Akaike weights to account for model selection uncertainty. Model assumptions and goodness of fit were examined for the best supported model using simulation-based residual diagnostics (e.g., dispersion, outliers, and residual uniformity) implemented in the *DHARMa* R package (Hartig 2016). Multicollinearity was further examined by calculating variation inflation factor using the *performance* R package (Lüdecke et al. 2021), which confirmed no serious collinearity among variables (all VIF values < 3; Zuur et al., 2010). A correlogram analysis based on Moran’s I indicated the absence of spatial autocorrelation in the model residuals (Fig. S6). The effect of the predictors was considered statistically significant based on Type II Wald tests (*p* < 0.05). To account for uncertainty associated with low-confidence sites, analyses were conducted both including these sites (n = 58) and excluding them (n = 50). The number of sites differed from that used in the elevational comparison analysis because habitat size could not be estimated for three sites. All statistical analyses were performed in R 4.5.1 (R Core Team 2025).

**Table 1.** Model selection results for the occupancy analysis ranked based on AICc showing the estimated predictor coefficients, the model degrees of freedom and the information criterion results with the corresponding model weights.

| Rank | Summer max temperature | Habitat size | Land use* | df | AICc | ΔAICc | Weight |
| --- | --- | --- | --- | --- | --- | --- | --- |
| 1 | -3.06 | 4.38 |  | 55 | 27.79 | – | 0.66 |
| 2 | -4.38 | 4.82 | + | 54 | 29.16 | 1.37 | 0.33 |
| 3 |  | 3.8 | + | 55 | 40.86 | 13.07 | 0 |
| 4 |  | 3.28 |  | 56 | 44.83 | 17.03 | 0 |
| 5 | -2.39 |  |  | 56 | 51.35 | 23.56 | 0 |
| 6 | -2.82 |  | + | 55 | 52.87 | 25.08 | 0 |
| 7 |  |  | + | 56 | 71.49 | 43.7 | 0 |
| 8 |  |  |  | 57 | 76.8 | 49 | 0 |

\* Inclusion of land use in the model is indicated by a plus sign (+) as it was a categorical variable (i.e., presence/absence of human land transformation within a 200-m radius centered at each site; see Methods for details).

## RESULTS

We detected presence of the northern pika at 41 sites out of the 61 sites surveyed (67.2%) (Figs 1, 2). This suggests that local populations of the northern pika in Hokkaido have become extirpated at 20 sites (32.8%). Population turnovers were observed at low-to mid-elevations, spanning a large elevational range of 60–1,547 m, with the median at 805 m (mean at 819 m) (Fig. 2). The lowest and highest absent sites in elevation were from the Samani-Erimo and Mt. Yubari regions, respectively. However, not all sites within this elevational range became absent, resulting in a sparser contemporary elevational distribution relative to the historical distribution (Fig. 2). At the regional level, we documented substantial variation in the proportion of present sites (Figs 1, S5). Whereas all 16 and 6 sites remained occupied (100 %) in the Mt. Tomuraushi and Mt. Hakuun regions, respectively, the proportion of present sites was 75.0 % (6/8 sites) in the Samani-Erimo region, 69.2% (9/13) in the Tokachi region, 9.1% (1/11) in the North Hidaka region, and 0% (0/4) in the Mt. Yubari region. This variation in the proportion of present sites did not show a strong association with the elevational gradient (Pearson’s *r* = 0.31, *p* = 0.55; Fig. S7). Unsurprisingly, the northern pika was absent in all sites lacking rock interstices, which corresponded with our identified low-confident sites (*n* = 8). Nonetheless, the elevational distribution pattern was generally consistent after removal of low-confident sites with 41 sites occupied out of the remaining 53 sites (77.4%), suggesting population turnover at 12 sites (22.6%) (Fig. S8).

**Fig. 2.**
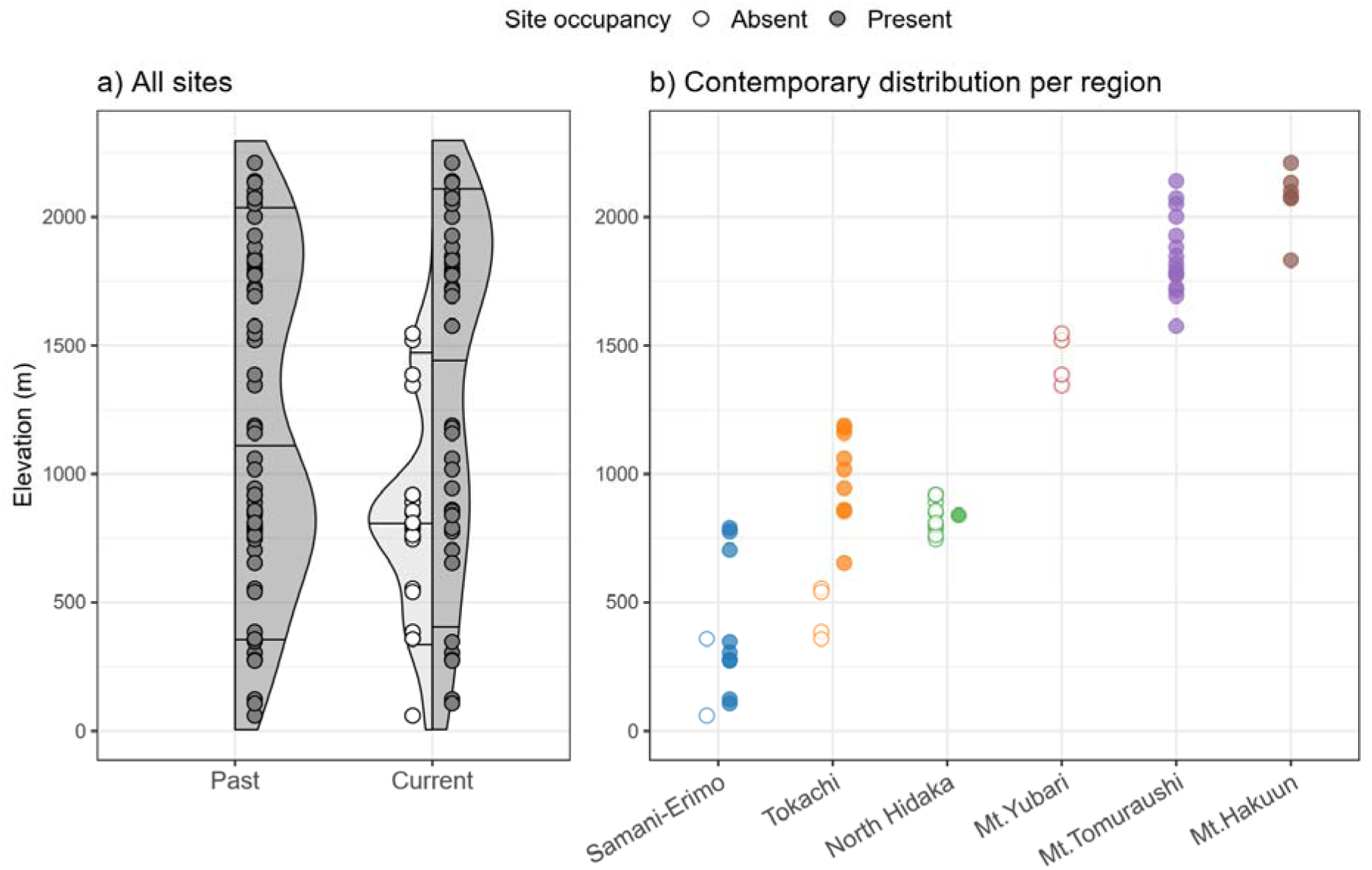
Elevational distribution of the northern pika, with presence and absence indicated by filled and unfilled points, respectively. (a) Comparison of elevational distribution between past and current surveys across all sites (*n* = 61). Shaded ribbons indicate frequency distribution of site elevations, with horizontal line segments indicating the 10th, 50th (median), 90th percentiles. (b) Elevational distributions of contemporary populations across regions.

Results from the quantile regression indicated a significant upshift in the central portion of the elevational distribution (50th percentile, median), from 1,018 m historically to 1,693 m in our resurvey (+675 m, *p* = 0.04) (Fig. 2). In contrast, elevational shifts were not significant at either the lower (10th percentile: -43 m, *p* = 0.84) or upper (90th percentile: +6 m, *p* = 0.93) portions of the distribution range. No statistically significant shift in mean elevation was also detected (two-sided *t*-test, *t* = -1.33, df = 100, *p* = 0.18). However, removing the low-confident sites turned the shift statistically nonsignificant for the central portion (50th percentile: +514 m, *p* = 0.14), while the lack of significance remained for the lower (10th percentile: 0 m) and upper (90th percentile: +6 m, *p* = 0.94) portions as well as the mean elevation (*t* = -0.94, df = 92, *p* = 0.35) (Fig. S8). These results suggest no clear evidence for a significant unidirectional shift after considering site location uncertainty.

In the occupancy analysis, habitat size and mean of summer maximum temperature were both included in the best-supported model, with one competing model (ΔAICc < 4) additionally including land use (Table 1). After model averaging, habitat size showed a significant positive effect (β = 4.33, *p* = 0.003) on persistence, whereas mean summer temperature had a significant negative effect (β = -2.13, *p* = 0.013) (Fig. 3). The effect of land use was nonsignificant (β = -0.10, *p* = 0.889). Because all predictors were standardized prior to analysis, our results indicate that habitat size exerted a stronger influence on persistence than mean summer temperature. These patterns were robust to the removal of low-confidence sites (Tables S3, S4).

**Fig. 3.**
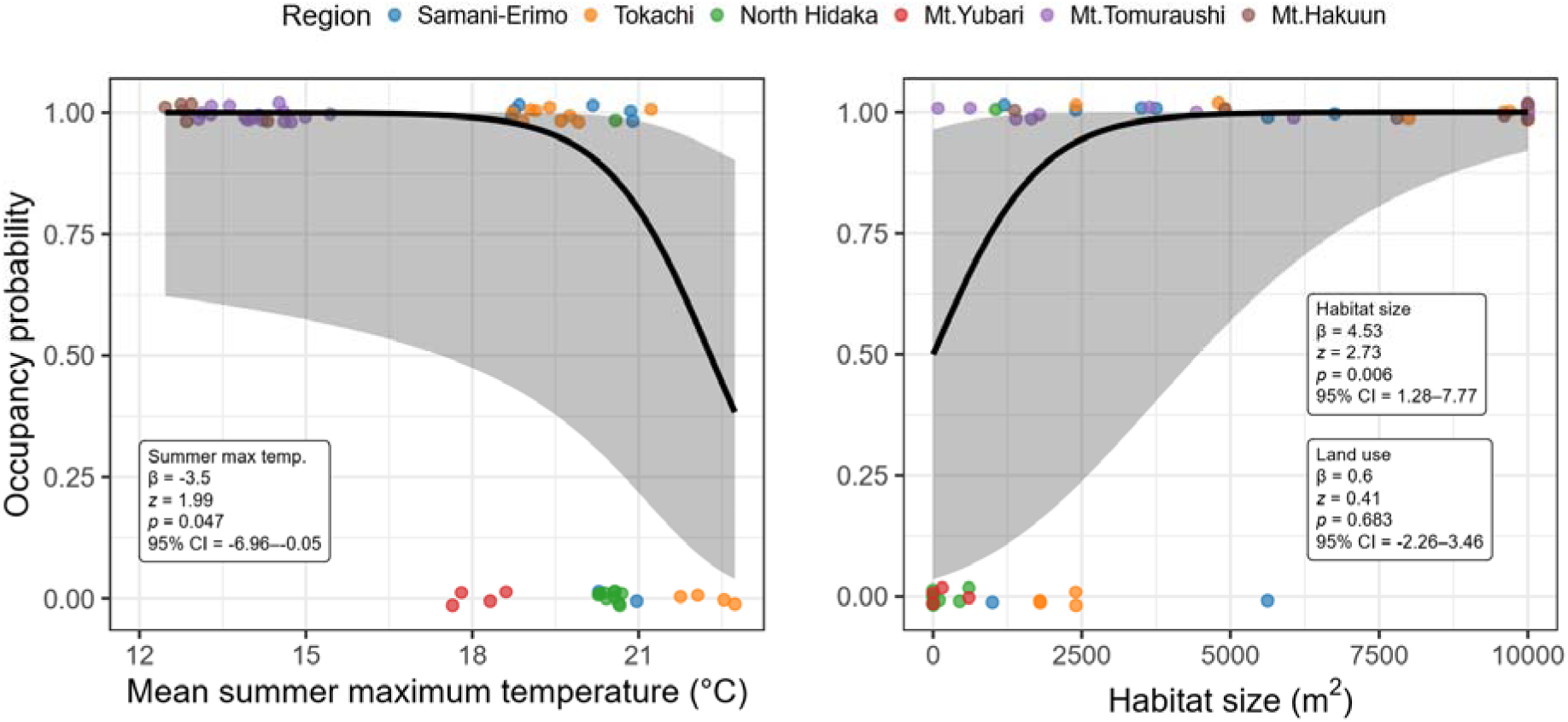
Predictions based on model-averaged coefficients in the GLM analysis using (left) mean of summer maximum temperature and (right) habitat size. Ribbons indicate 95% confidence intervals, and points represent raw data (1 = presence, 0 = absence) with jitter applied to avoid complete overlap. Although land use was included in the model as a categorical variable, both panels show predictions when land use is present. Predictions are similar when it is absent due to its minimal effect.

## DISCUSSION

Numerous resurvey studies have documented species distribution shifts along elevational gradients. The general focus in these studies is the link between temperature change and species distributions, while other climatic and environmental drivers are largely neglected (Rapacciuolo et al. 2014). Consequently, recent assessments on species redistribution are incorporating non-climatic factors to better capture the complex heterogeneous changes driving range shift responses (Lawlor et al. 2024). Here, we contribute to this growing body of evidence by documenting a possible ongoing shift in the elevational distribution of northern pika in Hokkaido from its historical distribution documented from the 1960s to the 2000s and its potential drivers. Over 30% of the historically occupied sites were currently vacant, all concentrated at low to mid elevations. Although these lower elevational landscapes are often prone to anthropogenic disturbances, our results show that temperature and habitat size alone explain the observed contemporary distribution and historical population turnover. This suggests that northern pika populations may be imperiled by higher temperatures at lower elevations even in the absence of human disturbance, while also highlighting the importance of larger habitats in facilitating persistence. Consequently, the remaining populations and their habitats represent a conservation priority that warrants careful monitoring.

We detected an upward shift in the central portion of the elevational distribution when all sites were included in the analysis, whereas shifts at the lower and upper portions remained statistically nonsignificant. However, after accounting for site location uncertainty by excluding low-confidence sites, the shift at the central portion also became nonsignificant. Taken together, these results suggest that current available evidence for an elevational shift exists although it is weak and remains unconclusive, highlighting the importance of continued long-term monitoring. Nonetheless, the observed pattern provides an opportunity to consider the processes that may lead to such a shift. Because each quantile represents a relative position within the overall elevational distribution, changes at the central portion should be interpreted in the context of where local populations disproportionately persisted or became extirpated. Upward expansions at leading distribution edges have been reported in other small mammals (Moritz et al. 2008). However, this was not detectable in our study because our dataset included only historically occupied sites. Consequently, the absence of an elevational shift at the upper portion of the distribution range was expected and likely reflects the suitable conditions existing at these higher elevation sites for the northern pika in Hokkaido (Sakiyama and García Molinos 2024, 2025). Moreover, the northern pika already inhabits the highest elevations of Hokkaido, leaving little opportunity for further upward expansion. In contrast, extirpations were more frequently observed from low-to mid-elevation sites, suggesting a contraction at the lower portion of the distribution, although this pattern was not statistically significant. This is possibly because extirpations occurred from low to mid elevations, while local populations persisted broadly across the entire elevational range. Consequently, despite declines in the number of occupied low-elevation sites, the relative position of the lower portion (10th percentile) remained comparatively stable. Under this interpretation, the upward shift detected at the central portion of the distribution may reflect the combined effects of persistence at higher elevations and partial extirpation at lower elevations, resulting in a greater relative concentration of occupied sites at higher elevations.

The concentration of extirpations in sites at low-to-mid elevations within the entire distribution suggests heat stress as the plausible trigger, which is supported by the negative effect of long-term mean of summer maximum temperature in the occupancy analysis. This finding agrees with previous studies documenting frequent absences of the northern pika at low to mid elevations in Hokkaido (Sakiyama et al. 2021, Sakiyama and García Molinos 2024), and similar studies of the American pika in North America (Rodhouse et al. 2010, 2018, Beever et al. 2011, Stewart et al. 2015, Billman et al. 2021). Pikas are generally considered vulnerable to higher temperatures because of their limited ability to dissipate heat (MacArthur and Wang 1973). Generally, increased physiological stress associated with higher temperatures is mitigated behaviorally, for instance, by using cooler areas available within their habitats and being active during the cooler times of the day. Although such responses are crucial for avoiding hyperthermia (i.e. behavioral thermoregulation; MacArthur & Wang, 1974; Onoyama, 1991; Smith, 1974; Smith et al., 2016), their effectiveness may have declined under ongoing climate warming in Hokkaido (Fig. S4), thereby affecting local population viability. Because a strong positive correlation was observed between the long-term mean and temporal change of summer maximum temperature, an alternative interpretation is that faster rates of climate warming may have contributed to extirpation, although the strong correlation between these variables prevented us from disentangling their individual effects.

Importantly, however, the observed variability in persistence (or extirpation) across sites at low to mid elevations suggests that such changes were not driven only by climate warming but also by other factors such as habitat size, which exhibited a strong positive effect on occupancy in our analysis. This result suggests that larger habitats facilitate persistence over time, possibly because larger habitats generally accommodate larger populations and have lower extinction risks resulting from demographic stochasticity (Griffen and Drake 2008). Moreover, larger habitats tend to exhibit greater environmental heterogeneity (Kohn and Walsh 1994), which may allow northern pikas to avoid predators more effectively and to utilize a wider range of forage resources than in smaller habitats. Rock interstice microclimates represent an important form of habitat heterogeneity that facilitates thermoregulation (Wilkening et al. 2015, Millar et al. 2016). Their greater availability and possibly variability in larger habitats may also have contributed to pika persistence under recent warming. Habitat size has been reported to be an important determinant of persistence in the American pika. Smith (1980) evidenced population stability in larger habitats at spatial scales matching those of our study although persistence was assessed over just five years; a much shorter time interval than our study. Moreover, different studies have showed the positive effect of larger habitats through resurveys carried out at temporal scales similar to ours (i.e. multi-decadal interval), while assessing habitat size at a larger spatial scale than our study (i.e., habitable area within a 1-km radius) (Beever et al. 2003, Stewart et al. 2015). This collective evidence suggests that habitat size drives the persistence of pikas ubiquitously across a wide range of temporal and spatial scales.

Presence of human land use did not significantly affect northern pika occupancy. This result contrasts with a previous study that detected a negative effect of land use (Sakiyama and García Molinos 2024), although this discrepancy may reflect differences in analytical scale, as the previous study focused only on lower elevation sites. Alternatively, the lack of a significant effect across the entire elevational range in our study may result from the simplified way in which the variable had to be estimated due to the lack of historical data. The binary classification based solely on current land use conditions did not account for historical changes in land use, nor did it capture the land use type and magnitude of land transformation. Future studies should therefore aim to overcome this oversimplification by integrating multiple sources of historical and spatial land use information where available.

Uncertainty in identifying the exact locations of previously reported sites posed a methodological challenge as some sites lacked rock interstices due to soil accumulation, litter, and vegetation cover (Fig. S3). Such uncertainty is a common issue in resurvey studies (Kapfer et al. 2017, Verheyen et al. 2017), particularly in vegetation research where sampling is conducted at small spatial scales at the meter level (Chytrý et al. 2014). In contrast, it is less likely that existing northern pika habitats were simply overlooked during field surveys given the distinctive rocky landforms and the spatial extent needed to support inhabitance. This pattern is also unlikely to be an artifact of cartographic inaccuracies in specific reports, as every report containing low-confidence sites also included sites where northern pika habitats could be successfully located. Instead, it is likely that the lack of suitable pika habitats at these low-confidence locations reflect actual environmental changes occurred at these sites over time rather than simple oversights in the previously reported locations. Indeed, based on descriptions in the original reports, we infer that these sites were not devoid of rock interstices at the time of historical surveys. For example, Haga et al. (1979) noted that “rocks were exposed” in the surveyed area while Kojima & Kawamichi (2001) explicitly reported the presence of “large rocks” and “rock interstices”. These descriptions contrast sharply with the current situation of these low-confident sites documented during our field observations (Fig. S3). One possible explanation is that these sites previously contained rock interstices abundant enough to support inhabitance, but that vegetation growth and soil accumulation over time eventually led to their loss, which accounts for an effective change in habitat size over time. However, this interpretation requires further investigation, as the factors driving large spatial variation in vegetation growth among nearby sites remain unclear. To our knowledge, vegetation dynamics within pika habitats have received limited attention despite their potential importance for understanding long-term persistence of local populations. Improved documentation of geographical locations and photographic evidence of habitat conditions would therefore be valuable in future studies.

We did our best to reduce the probability of misidentifying occupied sites by revisiting the study sites on multiple occasions and determine occupancy using playback, which is a highly effective detection method for this species (Sakiyama and García Molinos 2023). However, imperfect detections or false negative observations, cases where a species is deemed absent despite its presence, are inherently common in species distribution studies and can substantially influence inference (Bennett et al. 2024). Moreover, a recent study have shown that the American pika recolonized previously extirpated sites after more than 10 years (Millar and Smith 2022). Accordingly, we regard our study as an initial assessment of climate change impacts on the northern pika, and strong conclusions should not be drawn based solely on these results. Range contractions unfold as a succession of consecutive stages and related processes from an initial performance decline in existing populations followed by population decreases and their eventual local extinction (Bates et al. 2014). Although our results suggest substantial local extinctions, representing the final stage of this continuum, evidence from intermediate stages (e.g., declining populations at currently occupied sites) would provide an important extra layer of support for this interpretation. In this sense, while playback remains the most widely used method for surveying northern pika presence, the development of complementary and robust methodologies will be important for advancing our understanding of species behavior and population dynamics. For instance, there have been few attempts to utilize camera traps (Buchholz et al. 2021) and passive acoustic monitoring (Krause and Farina 2016) for studying the northern pika, both of which enable continuously, unmanned data collection and offer promising avenues for inferring behavioral patterns and estimating population dynamics.

Despite these limitations, our findings have important implications for predicting future changes and for the conservation of the northern pika in Hokkaido. As the potential threat from climate warming increases, the observed extirpation of numerous local populations may represent a precursor of a larger elevational contraction. Moreover, given the limited availability for upslope expansion as the species already occupy the available habitats at the upper end of its elevational distribution range, the northern pika in Hokkaido may experience a net reduction in its overall geographic range in the future, as predicted by a recent study (Sakiyama and García Molinos 2025), with potential consequences for the functioning of mountain ecosystems particularly in rocky landforms. Under such a scenario of latent extinction debts and considering the importance of habitat size evidenced by our results, the preservation and management of existing habitats across the range will become paramount to the future conservation of the species (e.g., Gheza et al., 2024). Accordingly, continuous monitoring of habitat conditions and local populations across the entire elevational distribution is urgently needed to detect further changes. However, there are currently no monitoring practices designed for this purpose for the northern pika despite its designation as Near Threatened in the Japanese Red List (Ministry of Environment 2020). We believe that the results from this study may contribute to developing a monitoring framework by adopting a repeatable methodology across the entire elevational distribution and detecting preliminary trends in the long-term persistence of local populations. Our results further highlight the critical importance of larger habitats for persistence of local populations, even under warmer conditions, whereas local populations in regions with few large habitats (i.e., Yubari and North Hidaka) appear particularly vulnerable. These results underscore the need to account for regional variability in population dynamics and to identify key habitats that can be prioritized for conservation at regional scales. Importantly, this study represents one of the few documented cases of species range shifts in terrestrial realms in Japan and highlights the need for similar assessments across other species.

In conclusion, this study represents the first attempt to explore the contemporary range shift of the northern pika in Hokkaido. Extirpations were detected at the central and lower portions of the distribution, with both climatic and non-climatic factors deemed likely causes for the observed changes. Future studies should increase resurvey sites and regions to better interpret the mechanisms underlying these changes. This study also calls for initiating practical conservation measures such as continuous monitoring programs to further detect changes in the overall distribution and monitor the viability of each regional population under climate change as well as protecting existing habitats.

## DATA AVAILABILITY

The data supporting the findings of this study are provided as Supplementary Material. The data and code used in this study will be deposited in a public repository upon publication of the associated peer-reviewed article.

## LITERATURE CITED

Bartoń, K. 2010. MuMIn: Multi-Model Inference:1.48.11.

Bates, A. E., G. T. Pecl, S. Frusher, A. J. Hobday, T. Wernberg, D. A. Smale, J. M. Sunday, N. A. Hill, N. K. Dulvy, R. K. Colwell, N. J. Holbrook, E. A. Fulton, D. Slawinski, M. Feng, G. J. Edgar, B. T. Radford, P. A. Thompson, and R. A. Watson. 2014. Defining and observing stages of climate-mediated range shifts in marine systems. Global Environmental Change 26:27–38.

Beever, E. A., P. F. Brussard, and J. Berger. 2003. Patterns Of Apparent Extirpation Among Isolated Populations Of Pikas (Ochotona Princeps) In The Great Basin. Journal of Mammalogy 84:37–54.

Beever, E. A., C. Ray, J. L. Wilkening, P. F. Brussard, and P. W. Mote. 2011. Contemporary climate change alters the pace and drivers of extinction. Global Change Biology 17:2054–2070.

Bennett, J. R., B. P. Edwards, J. N. Bergman, A. D. Binley, R. T. Buxton, D. E. Hanna, J. O. Hanson, E. J. Hudgins, S. Karimi, C. V. Raymond, C. D. Robichaud, and T. Rytwinski. 2024. How ignoring detection probability hurts biodiversity conservation. Frontiers in Ecology and the Environment 22:e2782.

Bergamini, A., S. Ungricht, and H. Hofmann. 2009. An elevational shift of cryophilous bryophytes in the last century – an effect of climate warming? Diversity and Distributions 15:871–879.

Bhatta, K. P., J.-A. Grytnes, and O. R. Vetaas. 2018. Downhill shift of alpine plant assemblages under contemporary climate and land-use changes. Ecosphere 9:e02084.

Billman, P. D., E. A. Beever, D. B. McWethy, L. L. Thurman, and K. C. Wilson. 2021. Factors influencing distributional shifts and abundance at the range core of a climate sensitive mammal. Global Change Biology 27:4498–4515.

Brook, B., N. Sodhi, and C. Bradshaw. 2008. Synergies among extinction drivers under global change. Trends in Ecology & Evolution 23:453–460.

Buchholz, R., J. Stamn, and S. A. Neha. 2021. Can camera traps be used to measure climate change induced alterations of the activity patterns of elusive terrestrial vertebrates? Climate Change Ecology 2:100020.

Burnham, K. P., and Anderson. 2002. Model Selection and Multimodel Inference. Springer, New Tork, NY.

Chen, I.-C., J. K. Hill, R. Ohlemüller, D. B. Roy, and C. D. Thomas. 2011. Rapid range shifts of species associated with high levels of climate warming. Science 333:1024–1026.

Chen, I.-C., S.-F. Shen, and S.-F. Chan. 2024. Niche Theory and Species Range Limits Along Elevational Gradients: Perspectives and Future Directions. Annual Review of Ecology, Evolution, and Systematics 55.

Chen, Y.-H., J. Lenoir, and I.-C. Chen. 2025. Limited evidence for range shift–driven extinction in mountain biota. Science 388:741–747.

Chytrý, M., L. Tichý, S. M. Hennekens, and J. H. J. Schaminée. 2014. Assessing vegetation change using vegetation-plot databases: a risky business. Applied Vegetation Science 17:32–41.

Crimmins, S. M., S. Z. Dobrowski, J. A. Greenberg, J. T. Abatzoglou, and A. R. Mynsberge. 2011. Changes in Climatic Water Balance Drive Downhill Shifts in Plant Species’ Optimum Elevations. Science 331:324–327.

Geospatial Information Authority of Japan. 2023. Digital Elevation Model. Geospatial Information Authority of Japan. (n.d.). Aerial images. https://service.gsi.go.jp/map-photos/app/.

Gheza, G., Z. Porro, M. Barcella, S. Assini, and J. Nascimbene. 2024. Habitat loss, extinction debt and climate change threaten terricolous lichens in lowland open dry habitats. Fungal Ecology 72:101384.

Gliwicz, J., J. Witczuk, and S. Pagacz. 2005. Spatial behaviour of the rock dwelling pika (*Ochotona hyperborea*). Journal of Zoology 267:113–120.

Gräler, B., E. Pebesma, and G. Heuvelink. 2016. Spatio-Temporal Interpolation using gstat. The R Journal 8:204–218.

Griffen, B. D., and J. M. Drake. 2008. A review of extinction in experimental populations. Journal of Animal Ecology 77:1274–1287.

Haga, R., Y. Fujimaki, and K. Onoyama. 1979. Mammals. Pages 1–24 in Hokkaido Government, editor. Survey Report of the Hidaka Mountain Ecosystem [Animals].

Hartig, F. 2016, August 26. DHARMa: Residual Diagnostics for Hierarchical (Multi-Level / Mixed) Regression Models.

Higashino, M., and H. G. Stefan. 2014. Hydro-climatic Change in Japan (1906-2005): Impacts of Global Warming and Urbanization. Air, Soil and Water Research 7:ASWR.S13632.

Kapfer, J., R. Hédl, G. Jurasinski, M. Kopecký, F. H. Schei, and J.-A. Grytnes. 2017. Resurveying historical vegetation data – opportunities and challenges. Applied Vegetation Science 20:164–171.

Kawabe, M. 2008. The distribution of *Ochotona hyperborea yesoensis* in Hokkaido. Bulletin of the Higashi Taisetsu Museum of Natural History 30:1–20.

Koenker, R. 2025. quantreg: Quantile Regression:6.1.

Koenker, R., V. Chernozhukov, X. He, and L. Peng, editors. 2017. Handbook of Quantile Regression. Chapman and Hall/CRC, New York.

Kohn, D. D., and D. M. Walsh. 1994. Plant Species Richness--The Effect of Island Size and Habitat Diversity. Journal of Ecology 82:367–377.

Kojima, N., and T. Kawamichi. 2001. A population study of Japanese pika (*Ochotona hyperborea*) in Mt. Yubari. Wildlife FORUM 6:149–154.

Krause, B., and A. Farina. 2016. Using ecoacoustic methods to survey the impacts of climate change on biodiversity. Biological Conservation 195:245–254.

Lawlor, J. A., L. Comte, G. Grenouillet, J. Lenoir, J. A. Baecher, R. M. W. J. Bandara, R. Bertrand, I.-C. Chen, S. E. Diamond, L. T. Lancaster, N. Moore, J. Murienne, B. F. Oliveira, G. T. Pecl, M. L. Pinsky, J. Rolland, M. Rubenstein, B. R. Scheffers, L. M. Thompson, B. Van Amerom, F. Villalobos, S. R. Weiskopf, and J. Sunday. 2024. Mechanisms, detection and impacts of species redistributions under climate change. Nature Reviews Earth & Environment.

Lenoir, J., R. Bertrand, L. Comte, L. Bourgeaud, T. Hattab, J. Murienne, and G. Grenouillet. 2020. Species better track climate warming in the oceans than on land. Nature Ecology & Evolution 4:1044–1059.

Lenoir, J., J. C. Gégout, P. A. Marquet, P. De Ruffray, and H. Brisse. 2008. A Significant Upward Shift in Plant Species Optimum Elevation During the 20th Century. Science 320:1768–1771.

Lenoir, J., and J.-C. Svenning. 2015. Climate-related range shifts - a global multidimensional synthesis and new research directions. Ecography 38:15–28.

Lüdecke, D., M. S. Ben-Shachar, I. Patil, P. Waggoner, and D. Makowski. 2021. performance: An R Package for Assessment, Comparison and Testing of Statistical Models. Journal of Open Source Software 6:3139.

MacArthur, R. A., and L. C. H. Wang. 1973. Physiology of thermoregulation in the pika, Ochotona princeps. Canadian Journal of Zoology 51.

MacArthur, R. A., and L. C. H. Wang. 1974. Behavioral thermoregulation in the pika *Ochotona princeps* : a field study using radiotelemetry. Canadian Journal of Zoology 52:353–358.

Mantyka-Pringle, C. S., P. Visconti, M. Di Marco, T. G. Martin, C. Rondinini, and J. R. Rhodes. 2015. Climate change modifies risk of global biodiversity loss due to land-cover change. Biological Conservation 187:103–111.

McCain, C. M., and J. Grytnes. 2010. Elevational Gradients in Species Richness. Page Encyclopedia of Life Sciences. First edition. Wiley.

Millar, C. I., and A. T. Smith. 2022. Return of the pika: American pikas re occupy long extirpated, warm locations. Ecology and Evolution 12.

Millar, C. I., R. D. Westfall, and D. L. Delany. 2016. Thermal components of American pika habitat—how does a small lagomorph encounter climate? Arctic, Antarctic, and Alpine Research 48:327–343.

Ministry of Environment. 2014. Red Data Book 2014.

Ministry of Environment. 2020. Ministry of Environment Red List 2020.

Morelli, T. L., C. Daly, S. Z. Dobrowski, D. M. Dulen, J. L. Ebersole, S. T. Jackson, J. D. Lundquist, C. I. Millar, S. P. Maher, W. B. Monahan, K. R. Nydick, K. T. Redmond, S. C. Sawyer, S. Stock, and S. R. Beissinger. 2016. Managing Climate Change Refugia for Climate Adaptation. PLOS ONE 11:e0159909.

Moritz, C., J. L. Patton, C. J. Conroy, J. L. Parra, G. C. White, and S. R. Beissinger. 2008. Impact of a century of climate change on small-mammal communities in Yosemite National Park, USA. Science 322:261–264.

Murakami, K. 2026, April 13. agrmesh. R.

Ohno, H., K. Sasaki, G. Ohara, and K. Nakazono. 2016. Development of grid square air temperature and precipitation data compiled from observed, forecasted, and climatic normal data. Climate in Biosphere 16:71–79.

Onoyama, K. 1991. Diurnal behaviors. Pages 56–65 *in* Hokkaido Government, editor. Survey Report of Wildlife Distribution: Pika Ecology.

Onoyama, K., T. Kurumada, and S. Ohmija. 1991. Ch.2 Home range. Pages 66–94 *in* Hokkaido Government, editor. Survey Report on Wildlife Distribution: Report on Pika Ecology.

Onoyama, K., and T. Miyazaki. 1991. Distribution in Hokkaido. Pages 25–55 *in* Hokkaido Government, editor. Survey Report on Wildlife Distribution: Survey Report of the Ecology of Pikas.

Parmesan, C., and G. Yohe. 2003. A globally coherent fingerprint of climate change impacts across natural systems. Nature 421:37–42.

Pebesma, E. J. 2004. Multivariable geostatistics in S: the gstat package. Computers & Geosciences 30:683–691.

Pecl, G. T., M. B. Araújo, J. D. Bell, J. Blanchard, T. C. Bonebrake, I.-C. Chen, T. D. Clark, R. K. Colwell, F. Danielsen, B. Evengård, L. Falconi, S. Ferrier, S. Frusher, R. A. Garcia, R. B. Griffis, A. J. Hobday, C. Janion-Scheepers, M. A. Jarzyna, S. Jennings, J. Lenoir, H. I. Linnetved, V. Y. Martin, P. C. McCormack, J. McDonald, N. J. Mitchell, T. Mustonen, J. M. Pandolfi, N. Pettorelli, E. Popova, S. A. Robinson, B. R. Scheffers, J. D. Shaw, C. J. B. Sorte, J. M. Strugnell, J. M. Sunday, M.-N. Tuanmu, A. Vergés, C. Villanueva, T. Wernberg, E. Wapstra, and S. E. Williams. 2017. Biodiversity redistribution under climate change: Impacts on ecosystems and human well-being. Science 355:eaai9214.

QGIS Development Team. 2023. QGIS Geographic Information System.

R Core Team. 2025. R: A Language and Environment for Statistical Computing. Vienna, Austria.

Rapacciuolo, G., S. P. Maher, A. C. Schneider, T. T. Hammond, M. D. Jabis, R. E. Walsh, K. J. Iknayan, G. K. Walden, M. F. Oldfather, D. D. Ackerly, and S. R. Beissinger. 2014. Beyond a warming fingerprint: individualistic biogeographic responses to heterogeneous climate change in California. Global Change Biology 20:2841–2855.

Rodhouse, T. J., E. A. Beever, L. K. Garrett, K. M. Irvine, M. R. Jeffress, M. Munts, and C. Ray. 2010. Distribution of American pikas in a low-elevation lava landscape: conservation implications from the range periphery. Journal of Mammalogy 91:1287–1299.

Rodhouse, T. J., M. R. Jeffress, K. R. Sherrill, S. R. Mohren, N. J. Nordensten, M. L. Magnuson, D. Schwalm, J. A. Castillo, M. Shinderman, and C. W. Epps. 2018. Geographical variation in the influence of habitat and climate on site occupancy turnover in American pika (*Ochotona princeps*). Diversity and Distributions 24:1506–1520.

Rowe, K. C., K. M. C. Rowe, M. W. Tingley, M. S. Koo, J. L. Patton, C. J. Conroy, J. D. Perrine, S. R. Beissinger, and C. Moritz. 2015. Spatially heterogeneous impact of climate change on small mammals of montane California. Proceedings of the Royal Society B: Biological Sciences 282:20141857.

Rowe, R. J., J. A. Finarelli, and E. A. Rickart. 2009. Range dynamics of small mammals along an elevational gradient over an 80-year interval. Global Change Biology:no-no.

Rubenstein, M. A., S. R. Weiskopf, R. Bertrand, S. L. Carter, L. Comte, M. J. Eaton, C. G. Johnson, J. Lenoir, A. J. Lynch, B. W. Miller, T. L. Morelli, M. A. Rodriguez, A. Terando, and L. M. Thompson. 2023. Climate change and the global redistribution of biodiversity: substantial variation in empirical support for expected range shifts. Environmental Evidence 12:7.

Sakiyama, T., and J. García Molinos. 2023. Efficacy of aural detection methods for detecting Northern Pika (*Ochotona hyperborea*) occupancy in rocky and densely vegetated habitats. Journal of Mammalogy.

Sakiyama, T., and J. García Molinos. 2024. Northern pikas experience reduced occupancy due to surrounding human land use despite the occurrence of suitable microclimates. Journal of Biogeography 51:1199–1212.

Sakiyama, T., and J. García Molinos. 2025. Mapping fine-scale distribution of the northern pika Ochotona hyperborea considering duality in microhabitat thermal conditions. Frontiers of Biogeography 18:e131541.

Sakiyama, T., and J. García Molinos. 2026, June 24. Dataset for: Thermal conditions and habitat size explain elevational range shift of the northern pika. Zenodo.

Sakiyama, T., J. Morimoto, O. Watanabe, N. Watanabe, and F. Nakamura. 2021. Occurrence of favorable local habitat conditions in an atypical landscape: Evidence of Japanese pika microrefugia. Global Ecology and Conservation 27:e01509.

Smith, A. T. 1974. The distribution and dispersal of pikas: influences of behavior and climate. Ecology 55:1368–1376.

Smith, A. T. 1980. Temporal Changes in Insular Populations of the Pika (Ochotona Princeps). Ecology 61:8–13.

Smith, A. T., C. H. Johnston, P. C. Alves, and K. Hackländer, editors. 2018. Lagomorphs: pikas, rabbits, and hares of the world. Johns Hopkins University Press, Baltimore.

Smith, A. T., J. D. Nagy, and C. I. Millar. 2016. Behavioral ecology of american pikas (*Ochotona princeps*) at Mono Craters, California: living on the edge. Western North American Naturalist 76:459.

Sokolova, N. A., I. A. Fufachev, D. Ehrich, V. G. Shtro, V. A. Sokolov, and A. A. Sokolov. 2024. Expansion of voles and retraction of lemmings over 60 years along a latitudinal gradient on Yamal Peninsula. Global Change Biology 30:e17161.

Stewart, J. A. E., J. D. Perrine, L. B. Nichols, J. H. Thorne, C. I. Millar, K. E. Goehring, C. P. Massing, and D. H. Wright. 2015. Revisiting the past to foretell the future: summer temperature and habitat area predict pika extirpations in California. Journal of Biogeography 42:880–890.

Tingley, M. W., and S. R. Beissinger. 2009. Detecting range shifts from historical species occurrences: new perspectives on old data. Trends in Ecology & Evolution 24:625–633.

Tingley, M. W., M. S. Koo, C. Moritz, A. C. Rush, and S. R. Beissinger. 2012. The push and pull of climate change causes heterogeneous shifts in avian elevational ranges. Global Change Biology 18:3279–3290.

Varner, J., and M. D. Dearing. 2014. The importance of biologically relevant microclimates in habitat suitability assessments. PLoS ONE 9:e104648.

Verheyen, K., P. De Frenne, L. Baeten, D. M. Waller, R. Hédl, M. P. Perring, H. Blondeel, J. Brunet, M. Chudomelová, G. Decocq, E. De Lombaerde, L. Depauw, T. Dirnböck, T. Durak, O. Eriksson, F. S. Gilliam, T. Heinken, S. Heinrichs, M. Hermy, B. Jaroszewicz, M. A. Jenkins, S. E. Johnson, K. J. Kirby, M. Kopecký, D. Landuyt, J. Lenoir, D. Li, M. Macek, S. L. Maes, F. Máliš, F. J. G. Mitchell, T. Naaf, G. Peterken, P. Petřík, K. Reczyńska, D. A. Rogers, F. Hø. Schei, W. Schmidt, T. Standovár, K. Świerkosz, K. Ujházy, H. Van Calster, M. Vellend, O. Vild, K. Woods, M. Wulf, and M. Bernhardt-Römermann. 2017. Combining Biodiversity Resurveys across Regions to Advance Global Change Research. BioScience 67:73–83.

Wang, X., D. Liang, W. Jin, M. Tang, Shalayiwu, S. Liu, and P. Zhang. 2020. Out of Tibet: genomic perspectives on the evolutionary history of extant pikas. Molecular Biology and Evolution 37:1577–1592.

Wilkening, J. L., C. Ray, and J. Varner. 2015. Relating sub-surface ice features to physiological stress in a climate sensitive mammal, the American Pika (*Ochotona princeps*). PLOS ONE 10:e0119327.

Wilson, R. J., D. Gutiérrez, J. Gutiérrez, D. Martínez, R. Agudo, and V. J. Monserrat. 2005. Changes to the elevational limits and extent of species ranges associated with climate change: Elevational shifts accompany climate change. Ecology Letters 8:1138–1146.

Zuur, A. F., E. N. Ieno, and C. S. Elphick. 2010. A protocol for data exploration to avoid common statistical problems: *Data exploration*. Methods in Ecology and Evolution 1:3–14.

