## Supporting Information for "Thermal conditions and habitat size explain elevational range shift of the northern pika"

**Table S1** Comparison of occupancy models fitted using mean annual temperature and mean summer maximum temperature over the last 44 years (1980–2023). The lower AICc value for the model using summer maximum temperature suggests that this variable better represents thermal effects on occupancy.

| Model structure | AICc | ΔAICc |
| --- | --- | --- |
| Occupancy ~  Summer maximum temperature | 51.4 | – |
| Occupancy ~  mean annual temperature | 67.3 | 15.9 |

**Table S2.** Correlation between continuous and categorical variables based on the point-biserial correlation test.

| Variable 1 | Variable 2 | r | *p* |
| --- | --- | --- | --- |
| Mean of summer maximum temperature | Human land use | 0.62 | < 0.001 |
| Habitat size | Human land use | -0.21 | 0.106 |

**Table S3.** Model selection result after removal of low confident sites.

| Rank | Summer max Temperature | Habitat size | Land use* | df | AIC | ΔAIC | Weight |
| --- | --- | --- | --- | --- | --- | --- | --- |
| 1 | -2.8 | 3.8 |  | 55 | 26.71 | 0 | 0.69 |
| 2 | -4.03 | 4.24 | + | 54 | 28.32 | 1.62 | 0.31 |
| 3 |  | 2.91 | + | 55 | 36.07 | 9.36 | 0.01 |
| 4 |  | 2.5 |  | 56 | 40.4 | 13.7 | 0 |
| 5 | -2.16 |  |  | 56 | 40.87 | 14.16 | 0 |
| 6 | -2.45 |  | + | 55 | 42.93 | 16.23 | 0 |
| 7 |  |  | + | 56 | 51.51 | 24.8 | 0 |
| 8 |  |  |  | 57 | 57.19 | 30.48 | 0 |

* Inclusion of land use in the model is indicated by a plus sign (+) as it was a categorical variable.

**Table S4.** Results of model-averaged coefficients after removal of low confident sites.

| Variable | Estimate | SE | z | p | 95% CI |
| --- | --- | --- | --- | --- | --- |
| Habitat size | 3.93 | 1.62 | 2.37 | 0.018 | 0.68 – 7.19 |
| Temperature | -3.18 | 1.64 | 1.89 | 0.059 | -6.47 – 0.12 |
| Land use [Present]* | 0.5 | 1.36 | 0.36 | 0.716 | -2.22 – 3.23 |

* As land use is a categorical variable, the estimate shown corresponds to the contrast between the level Present and the reference level Absent.

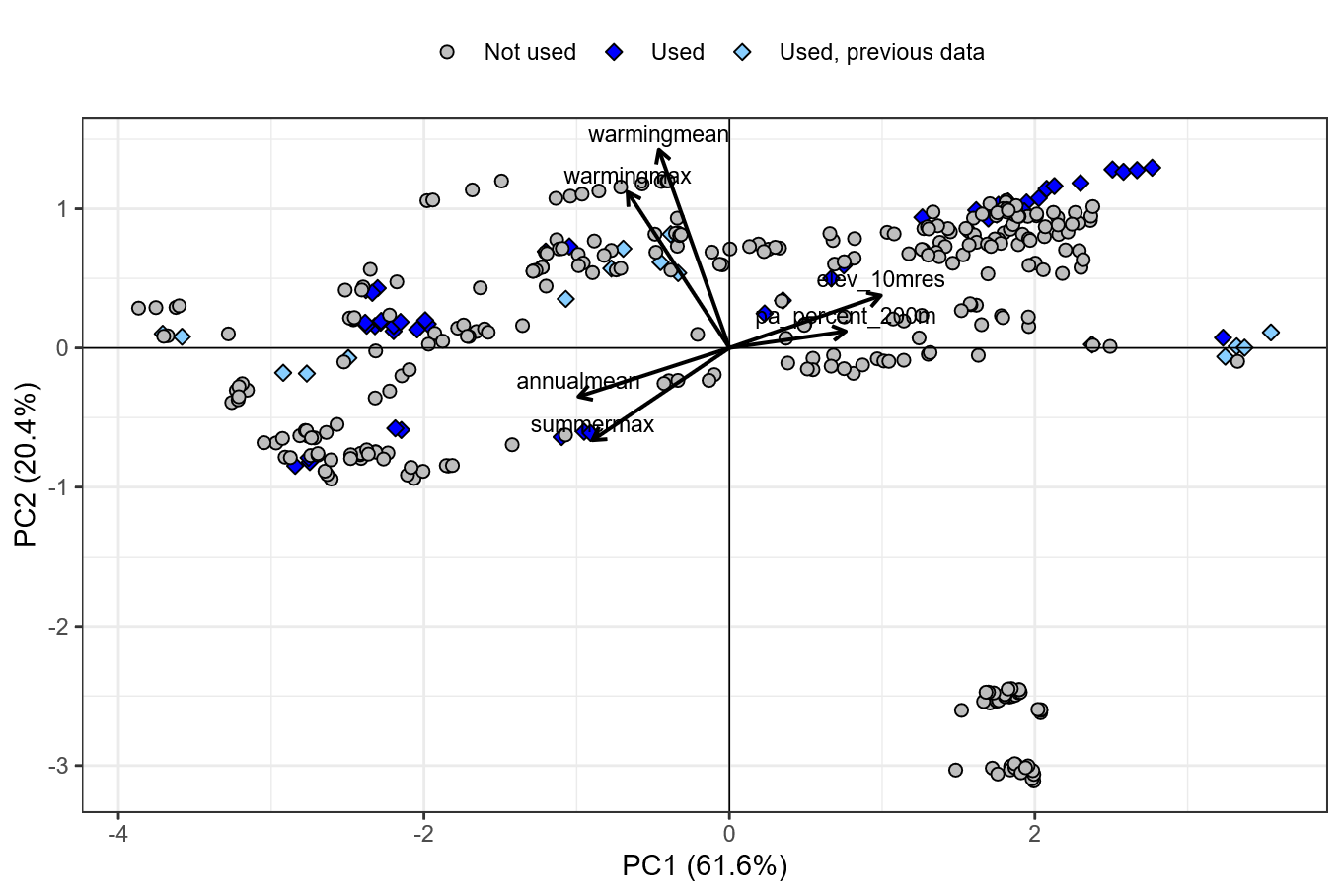

**Fig. S1** Result of the Principal Component Analysis conducted to select resurvey sites for this study. All points indicate sites where northern pika presence was detected in historical surveys. Particularly, site elevation, protection status (proportion of protected area within 200 m of the site), and thermal conditions, including mean and temporal change of mean annual temperature and summer maximum temperature over the past 44 years (1980–2023) were considered. The first two principal component axes (PC1 and PC2) together explained over 80% of the environmental variation among sites. Diamonds indicate sites used in this study (*n* = 61), whereas circles indicate sites not used in this study (*n* = 301). Among the diamond symbols, light blue indicates sites surveyed in a previous recent study (*n* = 16), whereas dark blue indicates sites newly surveyed in this study (*n* = 45).

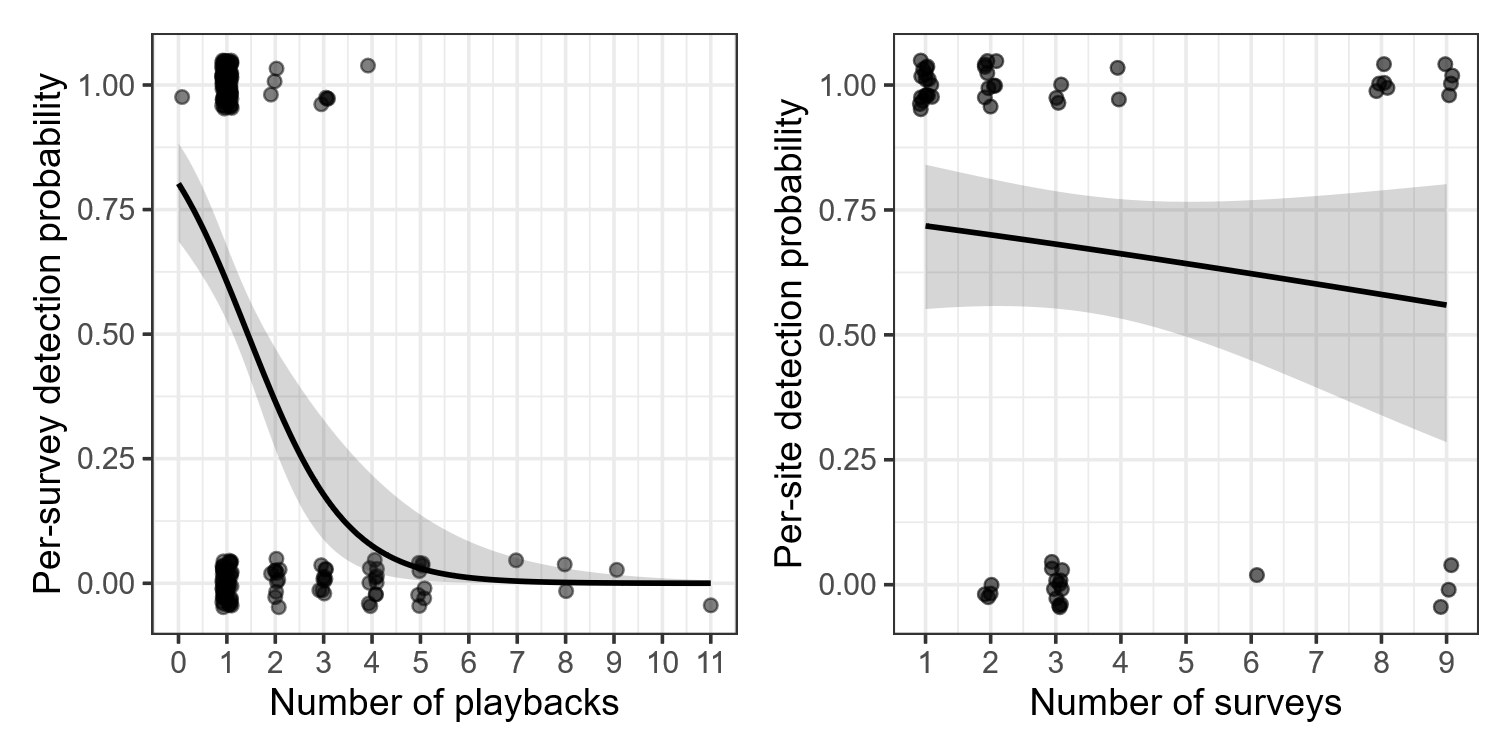

**Fig. S2** Response plots of logistic regression models examining detection probability per survey in relation to survey effort measured as the number of playbacks conducted (left), and detection probability per site in relation to survey effort measured as the number of surveys conducted (right). Per-survey detection probability was negatively associated with the number of playbacks (*β* = -0.98, *p* < 0.001), whereas no clear relationship was observed between per-site detection probability and the number of surveys conducted (*β* = -0.09, *p* = 0.37). These results do not support the commonly held assumption that greater survey effort increases detection probability. The negative relationship observed for per-survey detection may reflect the tendency for animal presence to be detected after only one or a few broadcasts, whereas repeated broadcasts were often conducted at sites where no animals were detected.

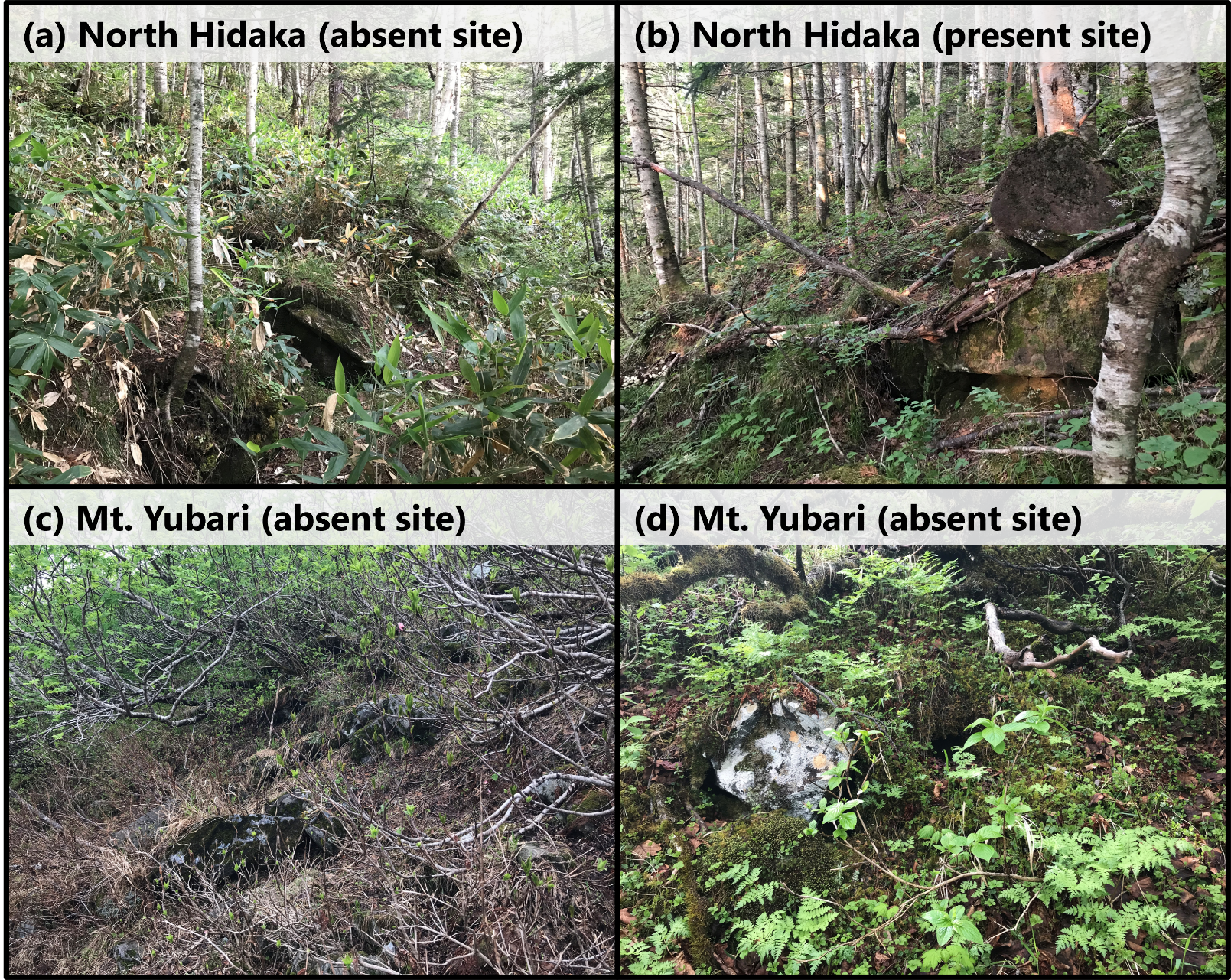

**Fig. S3.** Images of the resurvey sites. (a) A reported historically occupied site in the North Hidaka region lacking clear signs of rocky landforms due to soil formation, dense litter layer, and vegetation cover. Sites with such characteristics were identified as a low-confident sites for the purpose of the analysis (see Methods). The northern pika was not detected at these sites. (b) A site in the same North Hidaka region with sparse vegetation cover and clearly exposed rocky landforms, where the northern pika was detected. (c) Another low-confidence site in the Mt. Yubari region lacking clear signs of rocky landforms due to dense litter and vegetation cover. (d) Another location within the same site as (c), showing a small hole in the ground vegetation connected to rock interstices, which northern pikas commonly use for movement in densely vegetated habitats. However, such holes were limited in number, and the northern pika was not detected at this site.

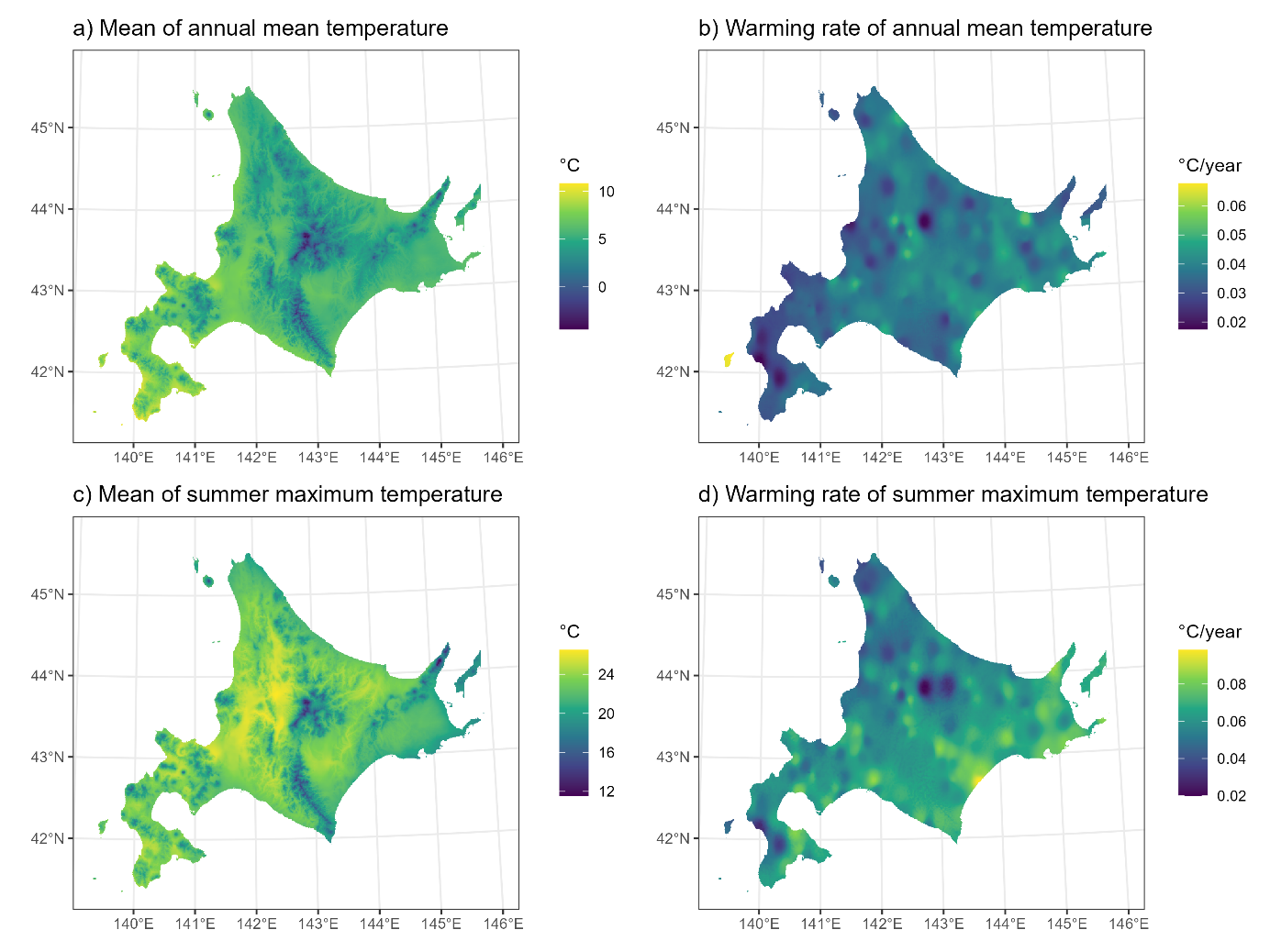

**Fig. S4** Temperature variables used in this study. Mean and temporal change in annual mean temperature and summer maximum temperature were calculated for the 44-year period (1980–2023). Panels correspond to (a) mean of annual mean temperature, (b) temporal change of annual mean temperature, (c) mean of summer maximum temperature, and (d) temporal change of summer maximum temperature. Raw temperature data were derived from the Agro-Meteorological Grid Square Data provided by the National Agriculture and Food Research Organization (see Methods).

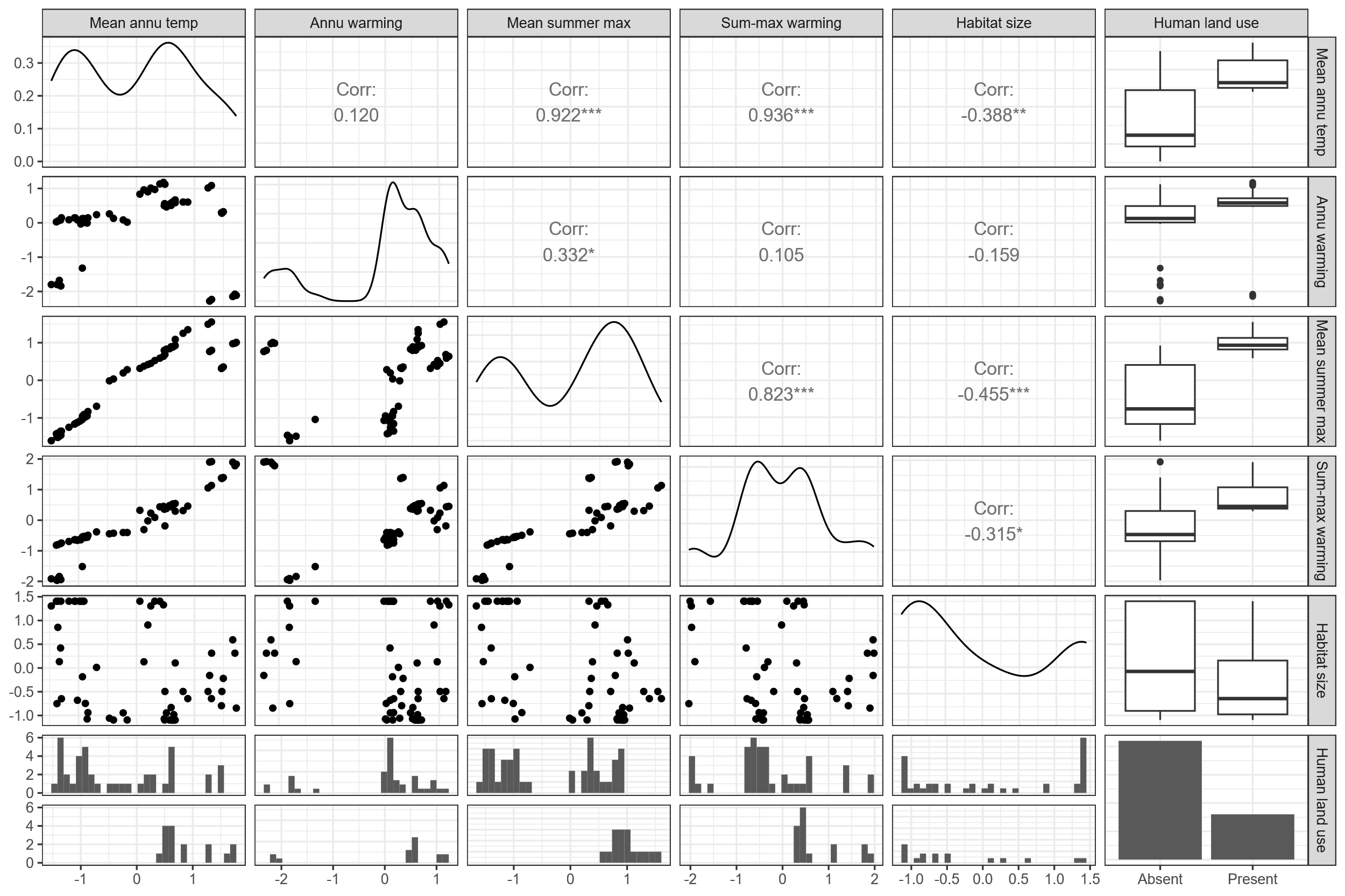

**Fig. S5.** Result of the pairwise correlation analysis among predictor variables. Variable name abbreviations correspond to: Mean annu temp. = Mean of annual mean temperature; Annu warming = Temporal change of annual mean temperature; Mean summer max = Mean of summer maximum temperature; Sum-max warming = Temporal change of summer maximum temperature.

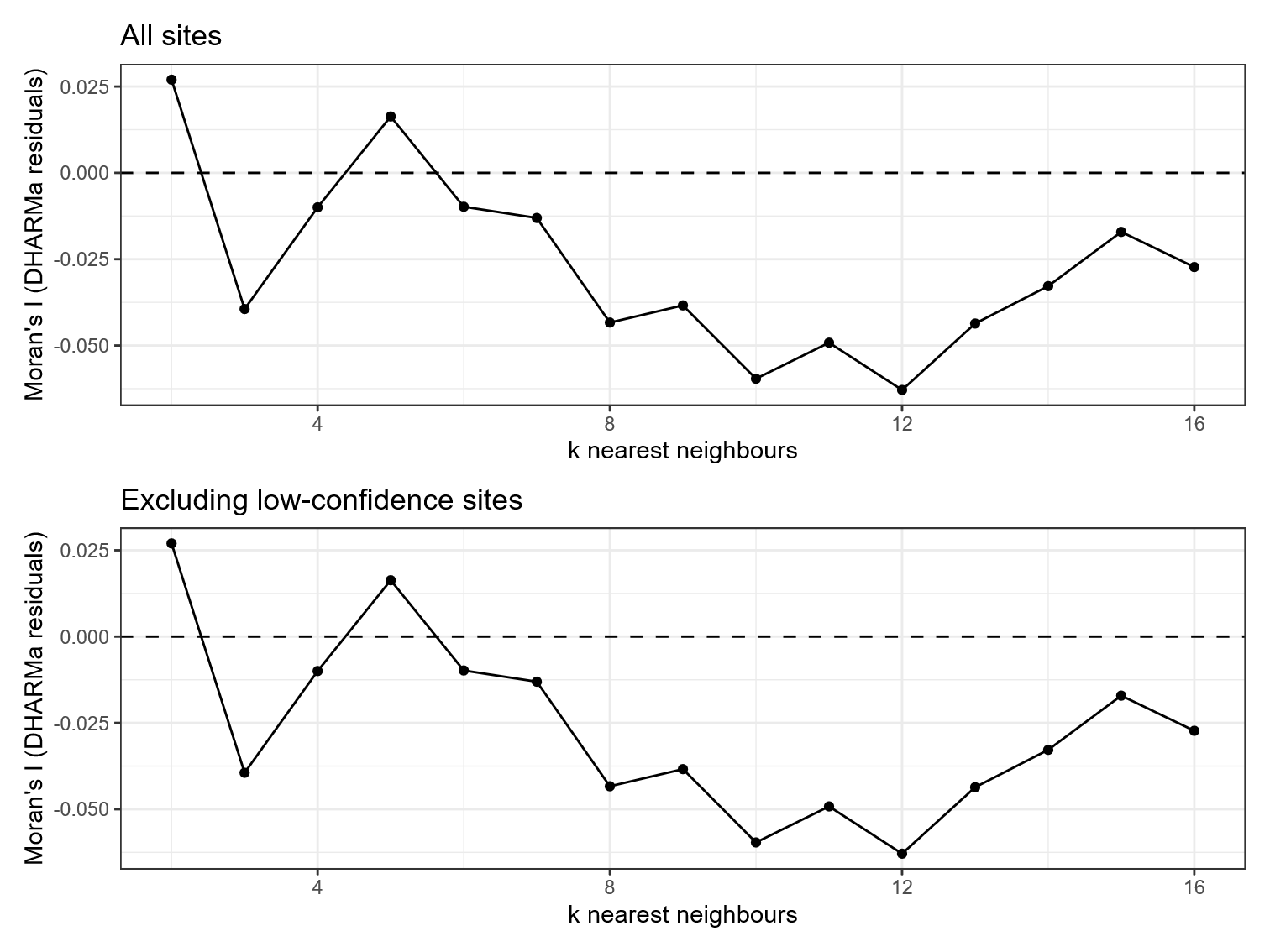

**Fig. S6.** Spatial correlogram of model residuals from the best supported generalized linear model for all sites (upper panel) and excluding low-confidence sites (lower panel).

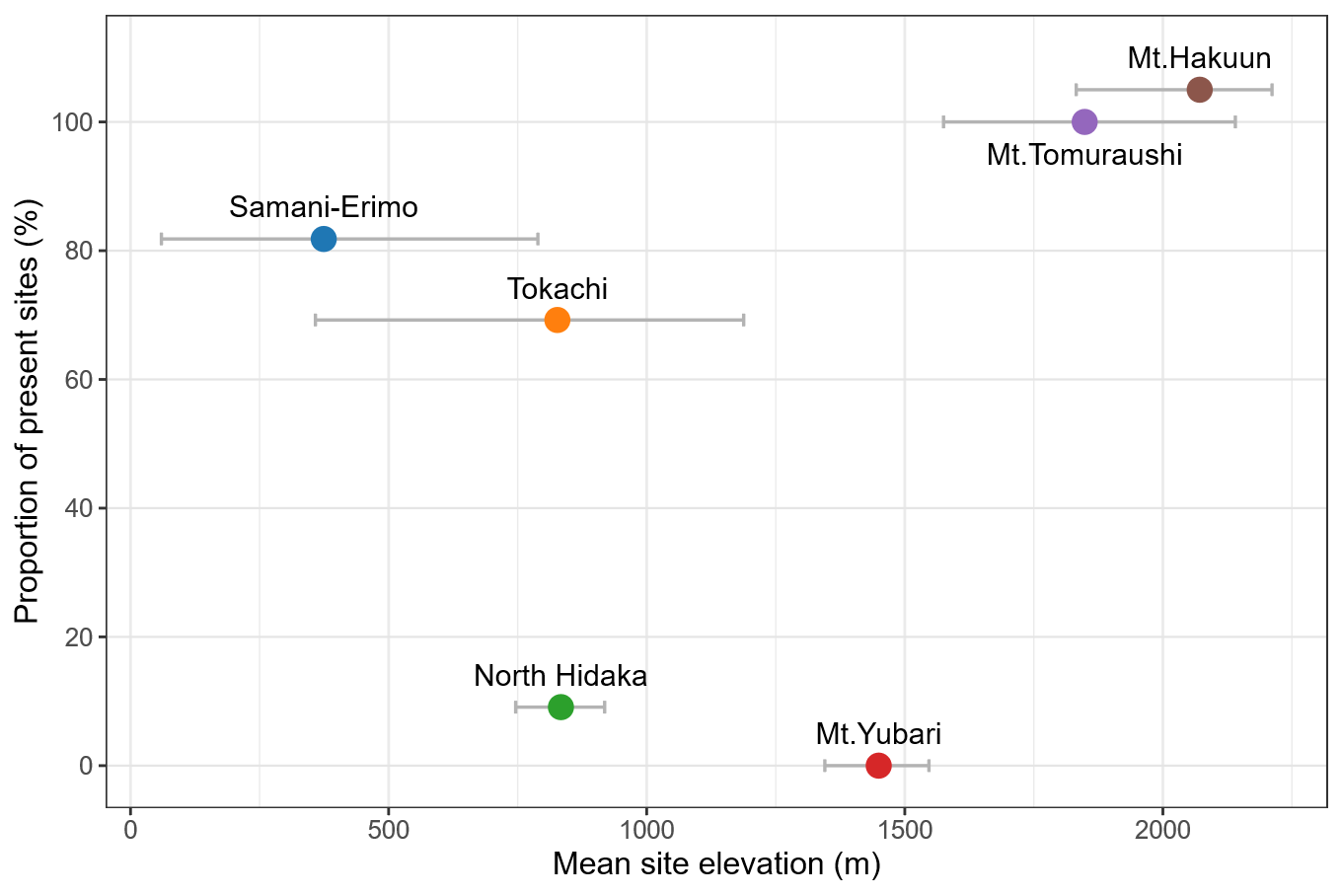

**Fig. S7.** Proportion of present sites plotted against mean site elevation within each region.

The horizontal bars denote the elevational range (minimum¬maximum) of sites within each region. The point and bar for Mt. Hakuun (100% persistence) were offset upward by 5 percentage points to prevent overlap with adjacent data.

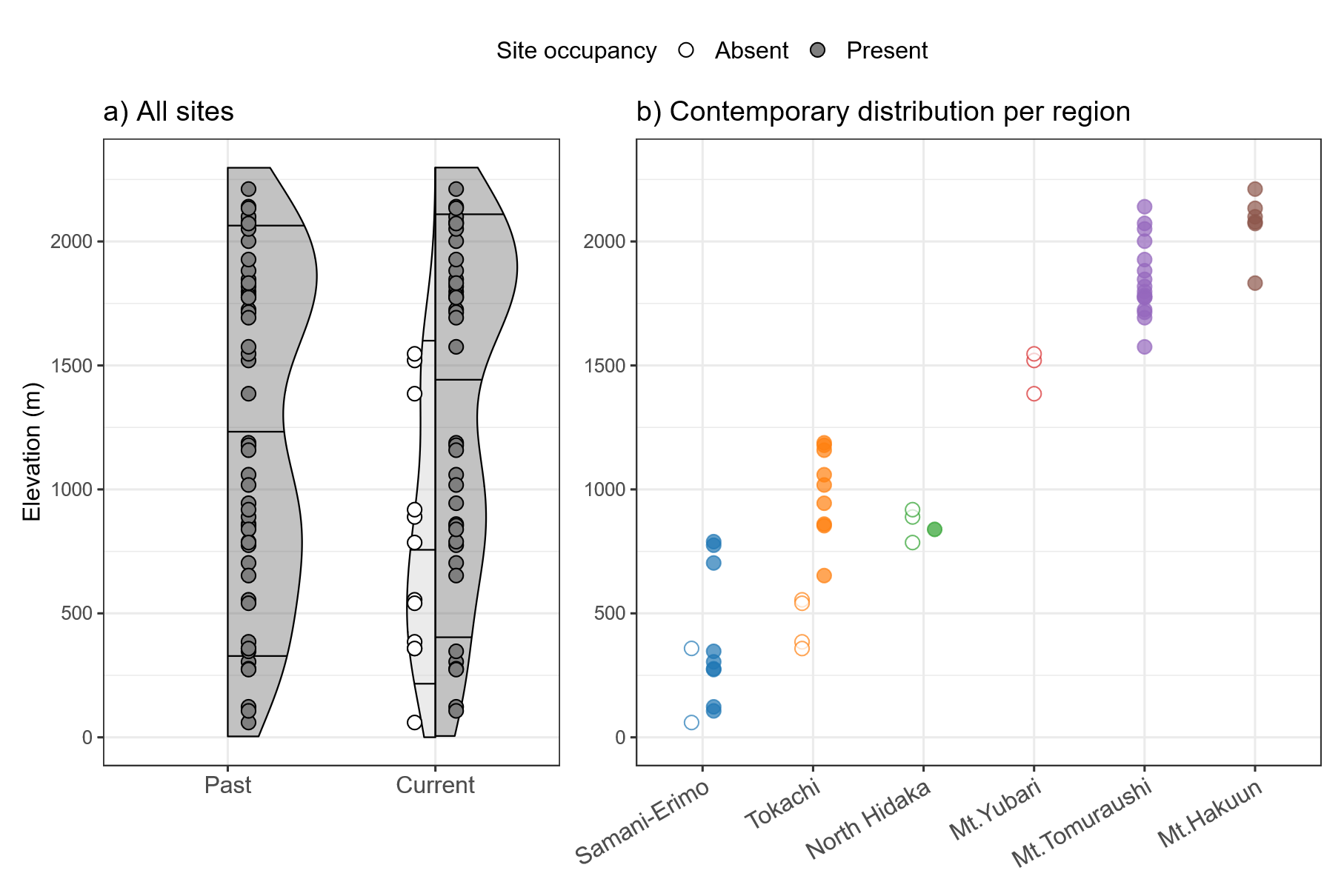

**Fig. S8.** Elevational distribution of the northern pika after removal of the low-confident sites, presence and absence indicated by filled and unfilled points, respectively. (a) Comparison of elevational distribution between past and current surveys across all sites. Shaded ribbons indicate frequency distribution of site elevations, with horizontal line segments indicating the 10th, 50th (median), 90th percentiles. (b) Elevational distributions of contemporary populations across regions.
